# Tycholyzer: NanoTemper NT.6 Tycho Analysis Tool

**DOI:** 10.64898/2026.11.13.766449

**Authors:** Vishnu Ganesh Ghani, Olga Boudker

## Abstract

NanoTemper Tycho NT.6 is a nano differential scanning fluorimetry (nanoDSF) instrument that enables rapid measurement of apparent protein melting temperatures. Thermal shift assays are widely used to assess protein stability under different conditions, including screening for protein-ligand interactions. These experiments often involve multiple samples, conditions, and replicates collected across several instrument runs, generating datasets that can be cumbersome to organize, compare, and analyze quantitatively. Here, we introduce Tycholyzer, a user-friendly software application that streamlines processing and analysis of Tycho nanoDSF data. Tycholyzer allows users to combine data from multiple runs, inspect and select individual fluorescence traces, and normalize and group replicate profiles for comparison across experimental conditions. The software provides two complementary approaches for determining apparent melting temperatures: sigmoidal fitting of fluorescence ratio profiles and identification of extrema in their first derivatives. Automated fitting of individual replicates enables calculation of mean melting temperatures and standard deviations, facilitating quantitative assessment of experimental variability. By integrating these capabilities into a single graphical interface, Tycholyzer promotes consistent, efficient, and rigorous analysis of nanoDSF experiments. Tycholyzer is freely available and can be run in MATLAB or as a standalone application using MATLAB Runtime (https://github.com/VGhani/Tycholyzer.git).

## 1 INTRODUCTION

Differential Scanning Fluorimetry (DSF) and nanoDSF are widely used techniques for monitoring protein unfolding^1^. DSF monitors fluorescence across a temperature gradient to identify one or more apparent protein melting temperatures (*T_M_*). Conventional DSF uses extrinsic labeling with fluorogenic dyes, such as SYPRO Orange, which increases its fluorescence upon binding to hydrophobic regions of the protein exposed upon unfolding^2^. Alternatively, the thiol-reactive dye 7-diethylamino-3-(4’-maleimidylphenyl)-4-methylcoumarin (CPM) becomes fluorescent upon reacting with cysteine residues exposed during unfolding^3^. As proteins unfold at rising temperatures, dye binding and fluorescence increase. Quantitative polymerase chain reaction instruments are often used to perform these assays.

In contrast, nanoDSF monitors the intrinsic fluorescence of aromatic residues, primarily tryptophan and, to a lesser extent, tyrosine, without requiring extrinsic dyes^4–6^. When excited at ∼280 nm, tryptophan residues are highly sensitive to their environment, emitting light between 310 and 380 nm. When buried in the hydrophobic protein interior, they exhibit emission maxima near 330-340 nm, whereas when exposed to an aqueous environment upon unfolding, their emission maximum shifts toward 350 nm^7,8^ The tryptophan fluorescence intensity and emission maximum can also be affected by proximity to neighboring amino acid side chains, including acidic and basic residues^9^. Consequently, protein thermal unfolding is often accompanied by changes in the intrinsic fluorescence spectra that can be monitored at 350 and 330 nm. The ratio of fluorescence intensities at these wavelengths provides a convenient measure of changes in the local environments of fluorescent residues and can be used to identify apparent thermal unfolding transitions^10,11^.

Thermal Shift Assays (TSAs) are widely used to screen for protein-ligand interactions^1,12,13^, optimize buffer conditions^12,14^, investigate the effects of lipid composition on membrane protein stability^12,15,16^, evaluate protein oligomerization states^17^, and analyze the effects of mutation on protein stability^18^. NanoTemper Tycho NT.6 (Tycho) performs nanoDSF measurements by exciting intrinsic protein fluorescence at 280-290 nm and measuring fluorescence intensities at 330 and 350 nm (*I_330_* and *I_350_*, respectively) during heating. The instrument applies a rapid temperature ramp from 35 to 95 °C at 30 °C/min, completing measurements in less than 3 minutes. Each Tycho run accommodates up to 6 capillaries and requires only ∼10 μl per sample at nanomolar concentrations, depending on the intrinsic protein fluorescence. Thermal unfolding is typically monitored using the temperature-dependent *I_350_*/*I_330_* ratio, and inflection temperatures, corresponding to *T_M_,* are determined from extrema in the first derivative of the ratio profiles.

Tycho exports data from each run into separate directories containing an Excel (xlsx) file with raw fluorescence data for all samples in the run. Experiments comparing multiple proteins, conditions, or replicates often require several Tycho runs. Consequently, the instrument generates multiple directories with output files that must be assembled for comparison and further analysis. Extracting and combining datasets from multiple runs for visual comparison, quality assessment, and quantitative analysis can be tedious and time-consuming. Although several software tools have been developed for DSF data analysis, including MoltenProt, TSA-CRAFT, and SimpleDSFviewer^10,19,20^, there remains a need for a streamlined workflow that facilitates the analysis and comparison of multiple Tycho datasets.

We developed Tycholyzer to address this need by integrating data management, visualization, and quantitative analysis into a user-friendly graphical interface. The software allows users to import data from multiple experimental runs, inspect and select individual fluorescence traces based on data quality, and automatically group and average replicates for comparison across experimental conditions. Tycholyzer provides two complementary approaches for determining apparent melting temperatures (Tₘ): sigmoidal fitting of fluorescence ratio profiles and identification of extrema in their first derivatives. Importantly, the software automatically fits individual replicate profiles and calculates mean Tₘ values and standard deviations, enabling quantitative assessment of experimental variability. Fitted values and summary statistics can be exported for further analysis. Together, these features simplify the analysis of large datasets and facilitate consistent, reproducible comparisons of protein thermal stability across experimental conditions.

## 2 IMPLEMENTATION

### 2.1 Overview and the workflow

Tycholyzer is a MATLAB-based application that enables users to organize, combine, and analyze traces from multiple xlsx files exported by Tycho. Users can run Tycholyzer in their native MATLAB workspace or by installing MATLAB Runtime on their system (see **Code Availability**). Tycholyzer imports multiple Tycho-generated xlsx files without modifying the original data and allows additional datasets to be incorporated into ongoing analyses. Software analysis sessions, including user selections and modifications, can be saved and subsequently reopened.

The Tycholyzer workflow consists of three main stages organized into five tabs (**Figure 1**). During the data import and quality assessment stage, the *Import Files* tab imports traces, enables visual assessment and selection of acceptable traces, and then collates and normalizes them. During the comparison stage, the *Inspect Sample Traces* tab visualizes individual and averaged replicate profiles to assess reproducibility and compare thermal unfolding behavior across experimental conditions. Finally, during the analysis stage, Tycholyzer supports two approaches. The *Sigmoidal Fit* tab fits sigmoidal curves to the *I_350_*/*I_330_* traces to obtain mean *T_M_* values with standard deviations, while the *Sigmoidal Fit (Advanced)* tab allows users to set initial guesses for the fitted parameters and the fitting temperature range. Alternatively, the *Inflection Point* tab obtains *T_M_* values from the first-derivative trace by identifying local extrema.

**Figure 1.**
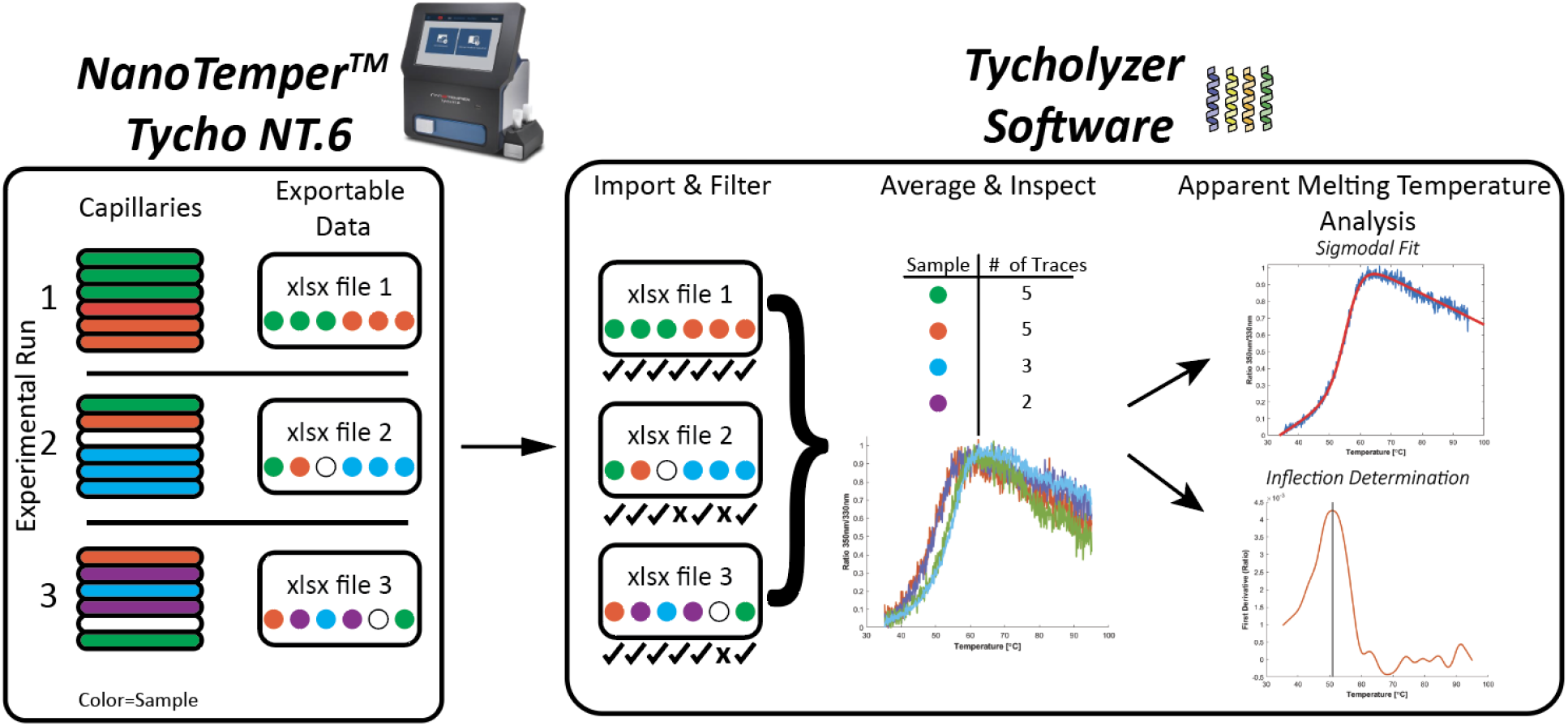
Experiment and analysis pipeline. In the Tycho instrument, users can load up to 6 samples per thermal melt run. Afterward, users can export experimental data as an xlsx file. In this example, three experimental runs are shown, with distinct samples/conditions represented by different colors and empty capillary lanes shown in white. After exporting files from the instrument, users import them into the Tycholyzer software. There, users can deselect traces unsuitable for analysis and group data for the same samples/conditions across all files for subsequent analysis to determine *T_M_* values.

The graphical interface allows users to move between data inspection, comparison, and analysis, and to swap visualizations of *I_350_*/*I_330_* ratio traces and their first derivatives using a toggle switch when screening and comparing data. A status lamp indicates whether the program is computing an action or is on standby. Finally, users can export graphs, tables, and numerical results for further analysis, documentation, and presentation.

### 2.2 Import Files tab

Users can import multiple Tycho data files into Tycholyzer simultaneously. The *Import Files* tab displays *I_350_/I_330_* traces (**Figure 2, left**) or first-derivative plots, controlled by a toggle switch. Users can inspect imported traces and exclude those with low signal-to-noise, optical interference, low-frequency noise, or other artifacts from further analysis. Poor data quality may result from low protein concentration, protein adsorption or precipitation, capillary imperfections, or interference from vibrations or condensation. Although initial sample *I_350_/I_330_* ratios may vary across replicates, the shape of the melting profile and corresponding *T_M_* values are often more consistent. To improve confidence in trace screening, comparison, and outlier identification, Tycholyzer normalizes each trace (**Section 4.2**) so that the minimum and maximum values are near 0 and 1, respectively (**Figure 2, right**). Following trace selection, users can press the “*Average selected data*” button to group accepted traces with identical capillary names across all imported files and calculate averaged normalized *I_350_/I_330_* and first-derivative profiles for subsequent analysis.

**Figure 2.**
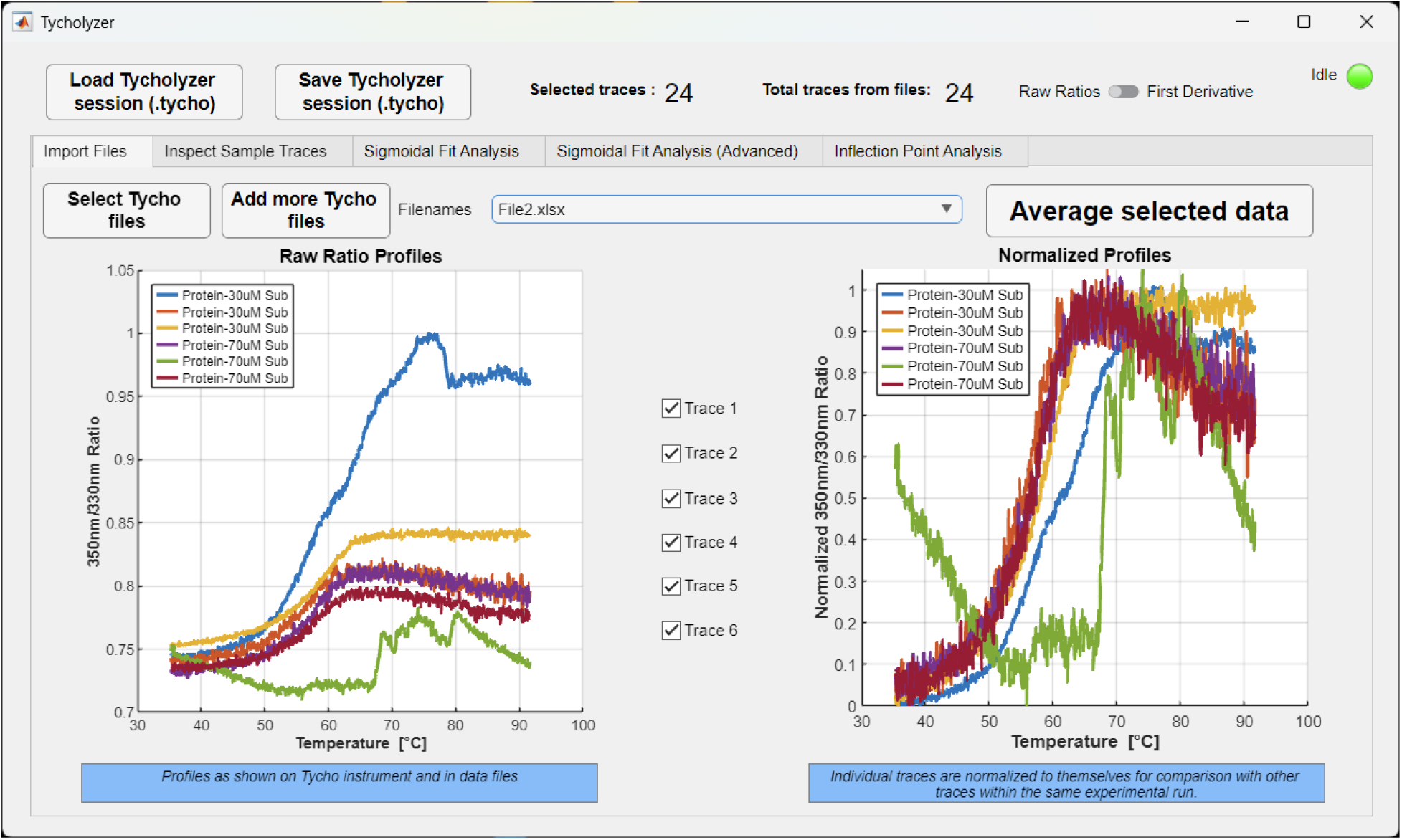
Import Files tab. The raw *I_350_/I_330_* ratio traces produced by Tycho and imported into Tycholyzer (left). Users can manually deselect undesirable capillaries that are empty or noisy, excluding them from further processing (e.g., blue and green traces). Because signals vary between capillaries, normalization (right) facilitates qualitative comparison of traces. For example, small differences in pre- or post-transition baselines (purple and red traces; orange and yellow traces) can produce seemingly distinct profiles. However, after normalization, they yield nearly identical melting profiles. After reviewing all files, users press the “Average selected data” button to group data across all imported files.

### 2.3 Inspect Sample Traces tab

After users select traces in the *Import Files* tab, pressing the “Average selected data” button populates the *Inspect Sample Traces* tab with the compiled data (**Figure 3**). Samples with identical names are automatically grouped, and the table lists distinct sample groups and the number of experimental traces in each. Users can manually rename samples to combine datasets that were assigned different names during data collection. Once renamed, Tycholyzer automatically combines the entries. These changes are applied within Tycholyzer without modifying the original excel files. To make the name change permanent, users have to edit the source xlsx file in the first worksheet named “Results”.

**Figure 3.**
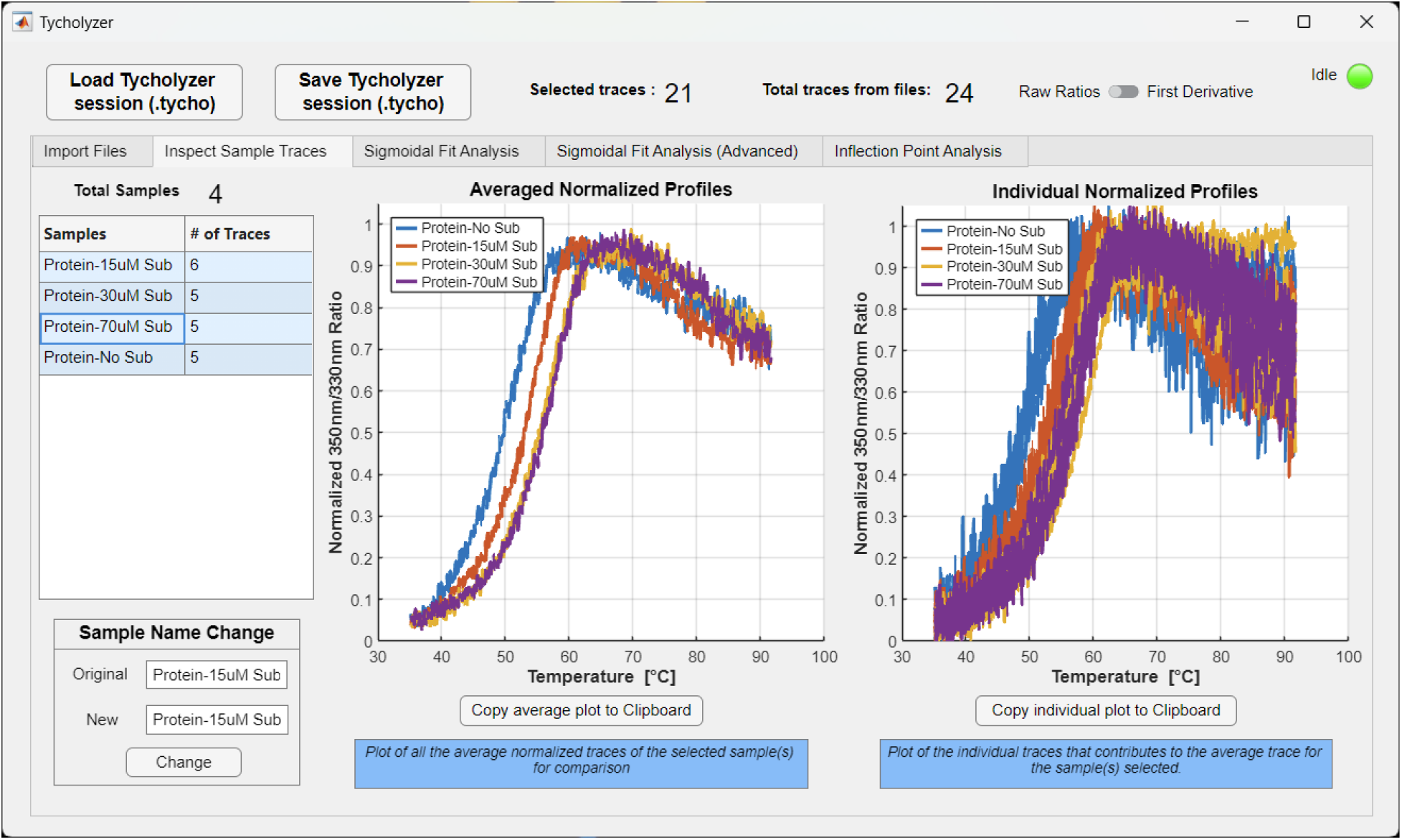
Inspect Sample Traces tab. After “Average selected data” compiles the screened data, the table in this tab populates with unique capillary names and the number of traces for each name. Selecting one or more sample group(s) from the table plots the average normalized trace for those samples (left) and the individual traces that contribute to the average (right). Selecting multiple sample groups enables qualitative comparison. Here, we compare four substrate concentrations and observe rightward shifts in the melting profiles as substrate concentration increases. In the corresponding individual plots, the same color-coded samples cluster within the sigmoidal transition region, while the post-transition baselines diverge more.

Selecting a sample group in the table displays its averaged normalized fluorescence ratio profile (**Figure 3, left**) and all individual traces that contribute to the average (**Figure 3, right**). Comparing individual traces helps users assess reproducibility, identify outliers that could skew the average, and determine whether additional replicates are needed. Multiple sample groups can also be selected simultaneously to qualitatively compare thermal unfolding profiles across all experimental conditions. Although Tycholyzer can accommodate an unlimited number of unique samples, a maximum of 12 traces can be displayed in a single plot. The generated plots can be copied directly into presentations or documents.

### 2.4 Sigmoidal Fit tab (Simple)

Selecting, normalizing, and averaging traces in the *Import Files* tab (**Figure 2**) also populates the *Sigmoidal Fit* tab (**Figure 4**). The “*Create Fits*” button in the tab fits the selected normalized fluorescence ratio profiles for each sample group to a sigmoidal function (**Section 4.3**) and returns the *T_M_* (**Figure 3**). Two complementary analyses are performed. First, the averaged normalized profile for each sample group is fitted to obtain a single *T_M_* value, reported in the left table. Users can select a sample group to display the averaged normalized trace (**Figure 3, blue**) and the fitted sigmoidal function (**Figure 3, red**). Second, each replicate trace is fitted independently, and the resulting *T_M_* values are used to calculate the mean and standard deviation for each sample group, reported in the right table. The *T_M_* values obtained by fitting the averaged profiles and by averaging the fitted *T_M_* values of individual traces are expected to be similar but not identical. Users can export both tables for further analysis and comparisons in their preferred program. The export option for the right table also includes the individual *T_M_* values.

**Figure 4.**
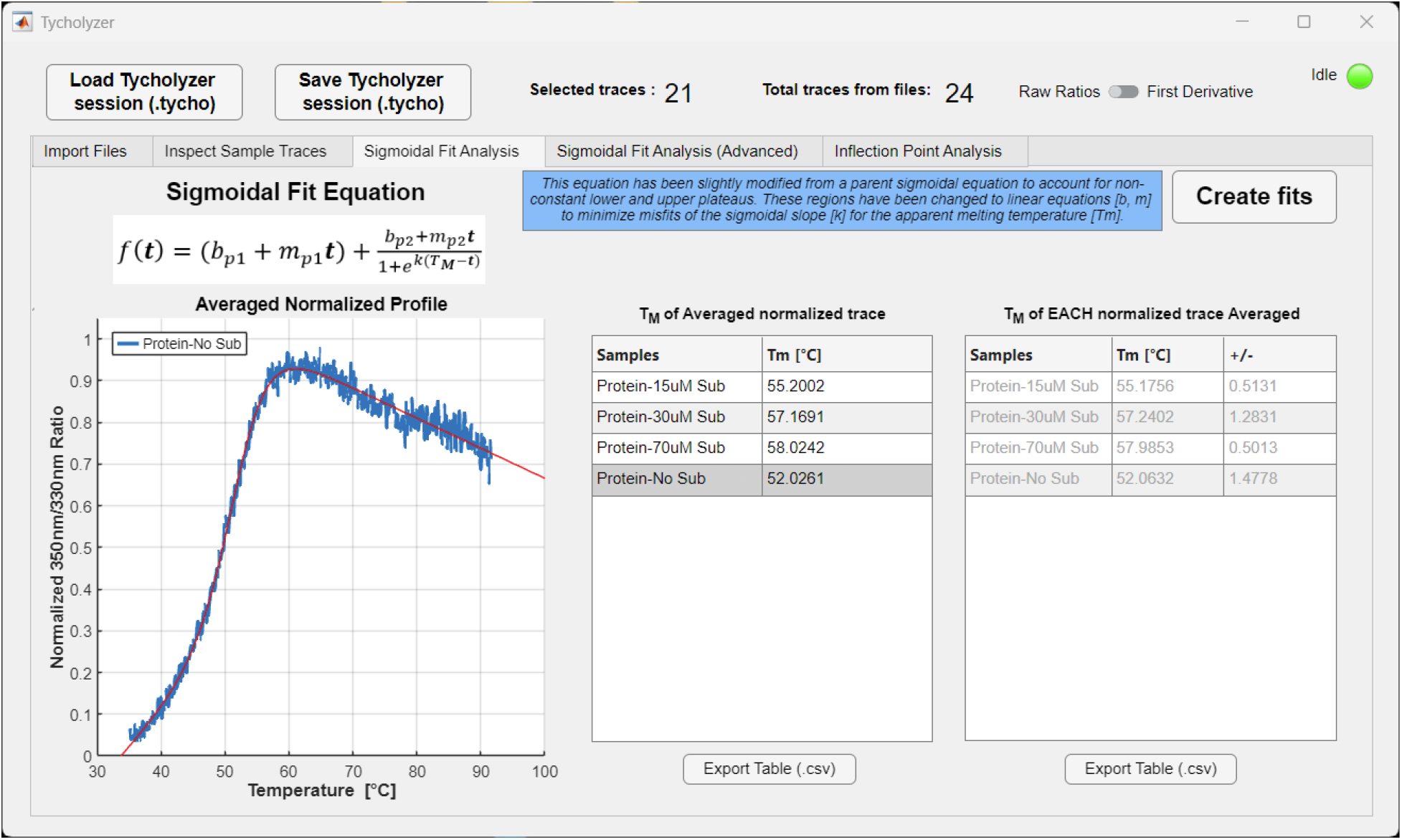
Sigmoidal Fit tab (Simple). When selected data is compiled using “Average selected data,” users can press the “Create fits” button to fit the data and obtain the fitted *T_M_* values for their sample group. The fits use a sigmoidal function modified to account for linear pre- and post-transition baselines. This function reports a single *T_M_* value and cannot account for multiple transitions. Two tables display melting temperatures determined by: (i) fitting the averaged normalized trace (left) and (ii) fitting the individual traces and averaging the resulting *T_M_* values for each sample group (right). By selecting a row in the left table, users can visualize the function fitted to the average normalized ratio trace.

If the sigmoidal fit fails, Tycholyzer displays an error message indicating the affected sample. Once the user acknowledges the message, the software continues fitting the remaining samples.

### 2.5 Sigmoidal Fit tab (Advanced)

*Sigmoidal Fit (Advanced)* provides additional control over the fitting procedure by allowing users to specify initial parameter guesses and restrict the temperature range used for fitting (**Figure 5**). These options are particularly useful when the default fitting procedure fails or produces unsatisfactory results, when multiple unfolding transitions need to be analyzed separately, or when artifacts in the pre- or post-transition baselines interfere with fitting. Users can first perform a basic fit, identical to that described in **Section 2.4**, to obtain initial estimates of the fitting parameters. These estimates can then be modified, and the corresponding “*Adv*” buttons activated to incorporate the user-specified values into the fitting procedure. Importantly, these values serve as initial guesses rather than fixed parameter values. Similarly, users can specify a temperature interval for fitting by activating the “*Adv*” option for *Fit Range*. Selecting “*Fit with Adv inputs*” performs the analysis using the specified settings. These settings are applied to all sample groups, rather than only the sample currently selected for display.

**Figure 5.**
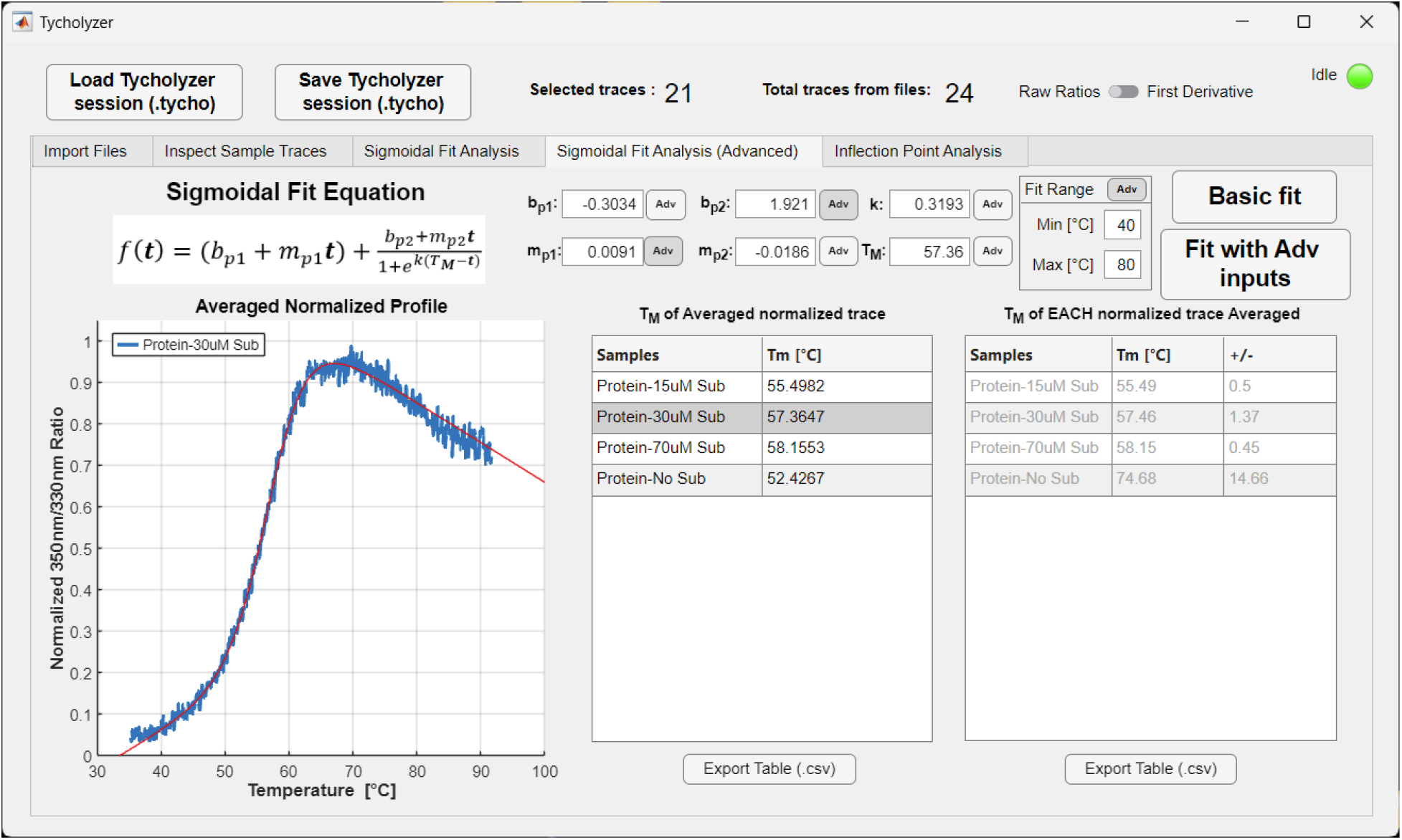
Sigmoidal Fit tab (Advanced). When selected data is compiled using “Average selected data,” users can press the “Basic fit” button to produce the same fitted T_M_ analysis as the simple Sigmoidal Fit tab. If fits are unsatisfactory or do not accurately parametrize the curve(s), users can adjust the initial coefficient guesses to improve them. Users will press the “Adv” button next to the module to switch to the advanced fit, then press “Fit with Adv inputs” to re-analyze all samples with the new settings. Users can also fit a selected range of temperatures. The fit will consider only the data within the range; however, the fitted curve will be plotted over the entire range of temperatures.

### 2.6 Inflection Point Analysis tab

The *Inflection Point Analysis* tab offers an alternative way to identify *T_M_* values from extrema in the first derivatives of fluorescence ratio profiles (**Figure 6**). After initiating the analysis with the “*Determine extrema*” button, users can select individual traces within each sample group to view their normalized first-derivatives profiles. Tycholyzer automatically identifies local minima and maxima (**Section 4.4**), marking their positions on the derivative plots and reporting the corresponding temperatures in the *Inflection Temp* table. This approach is similar to that used by the native Tycho software but uses a more sensitive algorithm that may detect additional extrema. Unlike sigmoidal fitting, derivative-based analysis can reveal multiple apparent unfolding transitions within a single profile. However, noise can also produce spurious extrema, including multiple peaks within a single transition. Users must therefore visually inspect the derivative profiles, select the extrema corresponding to meaningful thermal transitions, and manually record their temperatures. This approach provides a complementary method for analyzing complex unfolding profiles that may not be adequately described by a single sigmoidal function.

**Figure 6.**
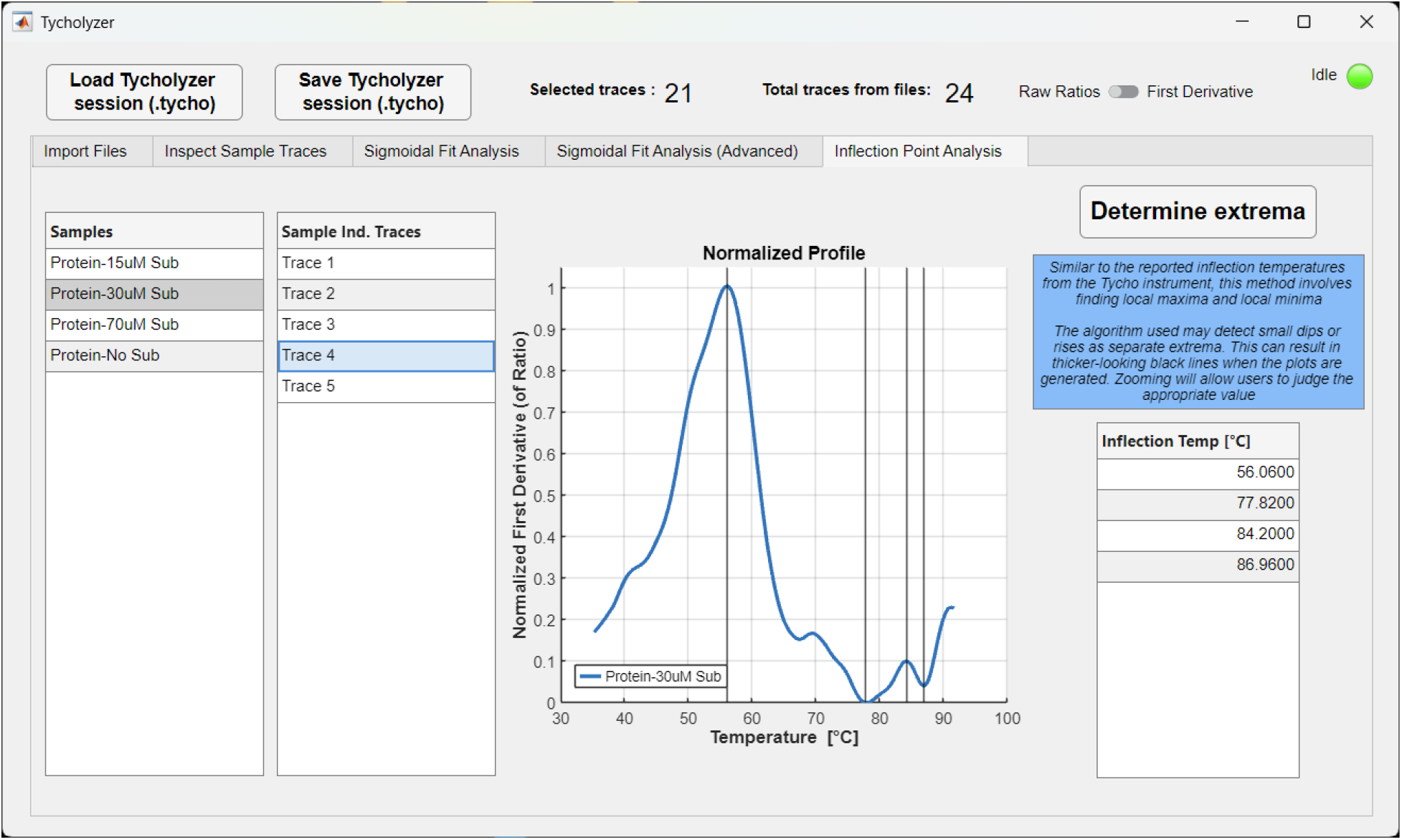
Inflection Point Analysis tab. After screening and compiling the data using the “Average selected data” button, users can click the “Determine extrema” button to populate the sample groups table (left table). Users can then select individual traces from the group (middle table) to visualize the normalized first-derivative plots showing the identified extrema. Users will then visually inspect the graph to determine whether the extrema are authentic and read the corresponding temperature in the Inflection Temp table (right table).

## 3 CONCLUSION

Tycholyzer provides a user-friendly, streamlined workflow for processing and analyzing nanoDSF data collected on the NanoTemper Tycho NT.6. The software facilitates compiling data from multiple experimental runs, assessing data quality, and comparing thermal unfolding profiles across samples, conditions, and replicates. Importantly, Tycholyzer enables automated fitting of individual and averaged fluorescence profiles to determine apparent melting temperatures and to calculate means and standard deviations across experimental replicates. An alternative approach based on the first derivatives of fluorescence profiles allows users to identify multiple thermal transitions. By integrating these capabilities into a single graphical interface, Tycholyzer reduces the time and effort required for data analysis and facilitates quantitative comparisons of protein thermal stability across experimental conditions.

## 4 DATA ANALYSIS ALGORITHMS

### 4.1 Data import from Tycho-generated Excel files

Tycholyzer reads raw data values and names from the input xlsx file(s). Each file contains four worksheets named Results, Profiles_raw, Profiles_smoothed, and Profiles_derivative. Capillary (sample) names are read from the Results worksheet in the capillary label column. Raw ratio data are read from the Profiles_raw worksheet, and first-derivative values of the ratio are read from the Profiles_derivative worksheet.

### 4.2 Trace normalization and averaging

Normalization is performed on both raw-ratio and first-derivative traces. For each trace, the lowest recorded value is subtracted from all data points, setting it to 0. Next, all data points are divided by the average of the resulting 15 maximum values.

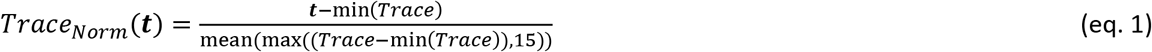

These normalized ratio traces and normalized first-derivative traces for each sample group are averaged for each recorded temperature value.

### 4.3 Sigmoidal fit of normalized ratio curves

*T_M_* values were obtained by fitting normalized individual or averaged traces to a modified sigmoidal function:

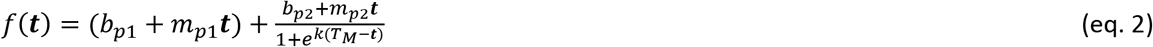

Where *b_pi_*, *m_pi_*: *i* = 1, 2 are, respectively, y-intercepts and slopes of pre- and post-transition baselines. *t* is the temperature value, *k* is the sigmoidal slope value, and *T_M_* is the melting temperature. Before non-linear curve fitting, normalized ratio traces are smoothed using a cubic spline with the MATLAB function csaps (p = .5). Smoothing is necessary to fit noisy traces. Erratic noise spikes, especially in the sigmoidal transition region, can greatly skew the fit or cause fitting failure.

### 4.4. Extrema detection for normalized first-derivative curves

Inflection points are determined by identifying extrema temperatures from the first-derivative profiles using the MATLAB function *findpeaks* with ’*MinPeakProminence*’ set to .05. Extrema values are detected with apparently higher sensitivity than by the software integrated into the Tycho instrument.

## CODE AVAILABILITY

Tycholyzer is a GUI developed in MATLAB App Designer and is publicly available (https://github.com/VGhani/Tycholyzer.git). The software is available for PC and Mac. Interested users can download and run it in MATLAB (v2023b or higher) or as a standalone application using the MATLAB Runtime (v2023b or higher). If running on a computer with MATLAB, users can download the *for_redistribution_files_only* directory and launch the software. If running on a computer without MATLAB, users can download the *for_redistribution* directory and run the item inside it. This will install the MATLAB Runtime (if not detected) and install the Tycholyzer software in the computer application directory.

## AKNOWLEDGEMENTS

We thank Krishna Reddy for encouraging the publication of this analysis software. We thank Poonam Dhankhar, Abdulkarim Farrokhzadeh, Farideh Badichi, Burha Rasool, and Alexander Earsley for testing preliminary software versions.

## AUTHOR CONTRIBUTIONS

V.G.G. conceptualized and designed the project, curated and analyzed the data, coded the software, and wrote the manuscript. V.G.G. and O.B. revised and edited the manuscript.

## ABRREAVIATIONS

DSF: Differential Scanning Fluorimetry
CPM: 7-diethylamino-3-(4’-maleimidylphenyl)-4-methylcoumarin
TSA: Thermal Shift Assay
T_M_: Apparent melting temperature Tycho - NanoTemper’s Tycho NT.6
Xlsx: Excel file exported by Tycho instrument
*I_350_*/*I_330_*: Ratio of intensities at 350 nm and 330 nm

